# Predicted Utility Modulates Working Memory Fidelity in the Brain

**DOI:** 10.1101/2021.04.01.438095

**Authors:** Emily J. Levin, James A. Brissenden, Alexander Fengler, David Badre

## Abstract

The predicted utility of information stored in working memory (WM) is hypothesized to influence the strategic allocation of WM resources. Prior work has shown that when information is prioritized, it is remembered with greater precision relative to other remembered items. However, these paradigms often complicate interpretation of the effects of predicted utility on item fidelity due to a concurrent memory load. Likewise, no fMRI studies have examined whether the predicted utility of an item modulates fidelity in the neural representation of items during the memory delay without a concurrent load. In the current study, we used fMRI to investigate whether predicted utility influences fidelity of WM representations in the brain. Using a generative model multivoxel analysis approach to estimate the quality of remembered representations across predicted utility conditions, we observed that items with greater predicted utility are maintained in memory with greater fidelity, even when they are the only item being maintained. Further, we found that this pattern follows a parametric relationship where more predicted utility corresponded to greater fidelity. These precision differences could not be accounted for based on a redistribution of resources among already-remembered items. Rather, we interpret these results in terms of a gating mechanism that allows for pre-allocation of resources based on predicted value alone. This evidence supports a theoretical distinction between resource allocation that occurs as a result of load and resource pre-allocation that occurs as a result of predicted utility.

## 1. Introduction

Working memory (WM) is central to human cognition. WM flexibly updates, temporarily maintains, and selects information to guide our behavior. It enables us to perform tasks effectively by both stably holding a goal in mind while resisting distractors, and flexibly updating that goal when the situation changes. However, WM is capacity-limited (Awh et al., 2007; Bays et al., 2009; Cowan, 2001; Luck & Vogel, 1997). As more information is maintained in memory, recall precision decreases (Bays et al., 2009; Wilken & Ma, 2004). This capacity limit fundamentally constrains the use of WM in service of cognition.

One way to make efficient use of this limited space is to prioritize relevant items depending on their utility. In other words, items that are more likely to be useful later are allocated more resources in WM and so are maintained either more robustly, with more precision, and/or for a longer duration.

Such prioritization is a feature of WM models that emphasize a gating control process for the management of WM (Braver & Cohen, 1999; Hazy et al., 2007; Hochreiter & Schmidhuber, 1997; Strock et al., 2020). In these models, a gating process selects items to be placed in WM, in a state robust to distraction, and to be selected from within WM as the context demands. Importantly, items are gated into WM to the degree that holding that item in memory is predicted to lead to future value. In other words, whether an item gets gated into WM or not depends on its predicted utility.

Prior evidence supports the overall hypothesis that predicted utility modulates the quality of information maintained in WM. Behavioral studies have shown that when items are prioritized based on their utility for an upcoming response, high-priority items are remembered with greater precision than low-priority items (Dube et al., 2017; Emrich et al., 2017; Fougnie et al., 2016; Gorgoraptis et al., 2011; Klyszejko et al., 2014; LaRocque et al., 2015; Ma et al., 2014; Yoo et al., 2018; Zokaei et al., 2011). Likewise, evidence from retro-cueing tasks scanned in fMRI has suggested that priority can modulate the fidelity of WM representations. In these tasks, participants must retroactively prioritize items that are already in memory based on a cue about which will be probed in an upcoming test. These studies have shown that fidelity of re-prioritized items is improved across a range of manipulations (Christophel et al., 2018; Larocque et al., 2017; Lewis-Peacock et al., 2012; Lorenc et al., 2020; Mallett & Lewis-Peacock, 2019; Sprague et al., 2016; Yu et al., 2020). Reframed in terms of predicted utility, the retrocue indicates which item, when maintained, is most likely to yield a positive outcome on that trial. Thus, together, these studies provide initial evidence that higher predicted utility results in higher-fidelity memory representations.

Importantly, however, it is not yet known whether items in WM are modulated as a function of their predicted utility independently of the effects of load. In retro-cueing studies, all items are initially encoded into memory and then a subset of those items is retroactively prioritized via a retro-cue. The retro-cued item is remembered with greater fidelity relative to the uncued items (Souza & Oberauer, 2016a). However, because the retro-cue is effectively reassigning predicted utility within a set of maintained items, it is not possible to distinguish whether changes to the quality of memory result from the predicted utility of that item or rather stem from a redistribution of a limited resource among items of different relative utility. In other words, can resources in memory be allocated based solely on the predicted utility of a target item itself, as opposed to only its relative utility to other items in memory?

One helpful clue to address this question comes from pre-cueing studies. In pre-cueing, a cue is presented before encoding that marks which items are likely to be tested. Pre-cues can range from being 100% valid, in which case cued items would have 100% predicted utility and uncued items would have 0% predicted utility, to having a continuous range of intermediate cue validities. In the latter case, all items must be remembered, but participants can still use the cue to prioritize items that are more likely to be probed. Importantly, this pre-cueing procedure allows for precision to be compared across different predicted utility states in a load-controlled way. Pre-cueing studies have shown that precision for remembered items significantly improves when items are cued beforehand (Bays, 2014; Bays & Husain, 2008; Dube et al., 2017; Emrich et al., 2017; Zokaei et al., 2011). This is important for the current study because it suggests that WM precision is influenced by factors other than load, and it provides additional evidence that WM resources can be strategically allocated to remembering prioritized items regardless of their utility relative to other items held concurrently. However, evidence of these graded effects is exclusively behavioral. Prior pre-cued WM fMRI studies have used 100% valid cues that, accordingly, have 100% predicted utility or 0%, if they are uncued (McNab & Klingberg, 2008). No fMRI studies to date have looked at how WM fidelity changes with parametrically increasing graded levels of predicted utilities in a load-controlled way. Thus, it remains an open question whether people control the fidelity of an item held in WM as a function of its predicted utility throughout a delay and independently from load.

In the current study, we sought to test the hypothesis that predicted utility influences the fidelity of WM representations in the brain continuously and independently of the influence of load. We had participants perform a modified WM gating task (the context-first context-last) (Chatham et al., 2014) adapted for continuous report in an fMRI scanner to test how predicted utility affects high-fidelity visual codes in WM. Across conditions of the task, a context cue determined whether the first oriented grating encountered was predicted to have high (100%), medium (50%), or no (0%) utility for the upcoming continuous report test. Using a generative model approach (Brissenden et al., 2021; van Bergen et al., 2015), we reconstructed from visual cortical fMRI activity the orientation stored in WM, as well as the uncertainty with which that orientation was encoded. To preview our results, we do find evidence that the fidelity of the remembered orientation is modulated by the predicted utility of the grating such that lower utility items are remembered with lower fidelity, even when they are the only item being maintained.

## 2. Methods

### 2.1 Participants

11 participants were recruited for this experiment. All participants were right-handed, had normal or corrected-to-normal vision, and had no history of neurological or psychiatric disorders. Two participants withdrew prior to completion of the first scanning session. A third participant chose not to return for the second session. A fourth participant completed both sessions but was excluded prior to analysis due to movement greater than our voxel size (2.4 mm). Thus, 7 participants were included in the study (5 male; mean age: 27.1 years; mean years of education: 17.6 years). These participants each completed a 1-hour behavioral training session in which they learned and practiced the cognitive task, and two 2-hour fMRI scanning sessions. The sample in this multi-session study was comparable to that in other multivariate encoding and decoding working memory studies (Brouwer & Heeger, 2009, 2011; Ester et al., 2015; Harrison & Tong, 2009; Lorenc et al., 2020; Pratte & Tong, 2014; Sprague et al., 2014, 2016; Sprague & Serences, 2013), and favors collecting more data across sessions in fewer participants in order to address significant between-participant variability. Participants were compensated $20/hour for fMRI sessions and $10/hour for behavioral training sessions. All participants gave written informed consent to participate in the experiment. The study was approved by the Research Protections Office at Brown University.

### 2.2 Experimental design

#### 2.2.1 Logic and Design

We designed a continuous report orientation task (Fig. 1) that allowed us to manipulate predicted utility and measure the resulting effects on precision. This task was modified from a 2AFC working memory task previously developed by our laboratory (Chatham et al., 2014) into a continuous report paradigm that required participants to maintain a high-fidelity visual code in working memory. Participants were presented with two orientation gratings on every trial. Depending on the type of trial, they had to remember one or both gratings. At the end of every trial, we tested participants’ working memory precision by having them rotate a randomly oriented response wheel so that it matched the angle of the target grating they were holding in memory.

**Figure 1.**
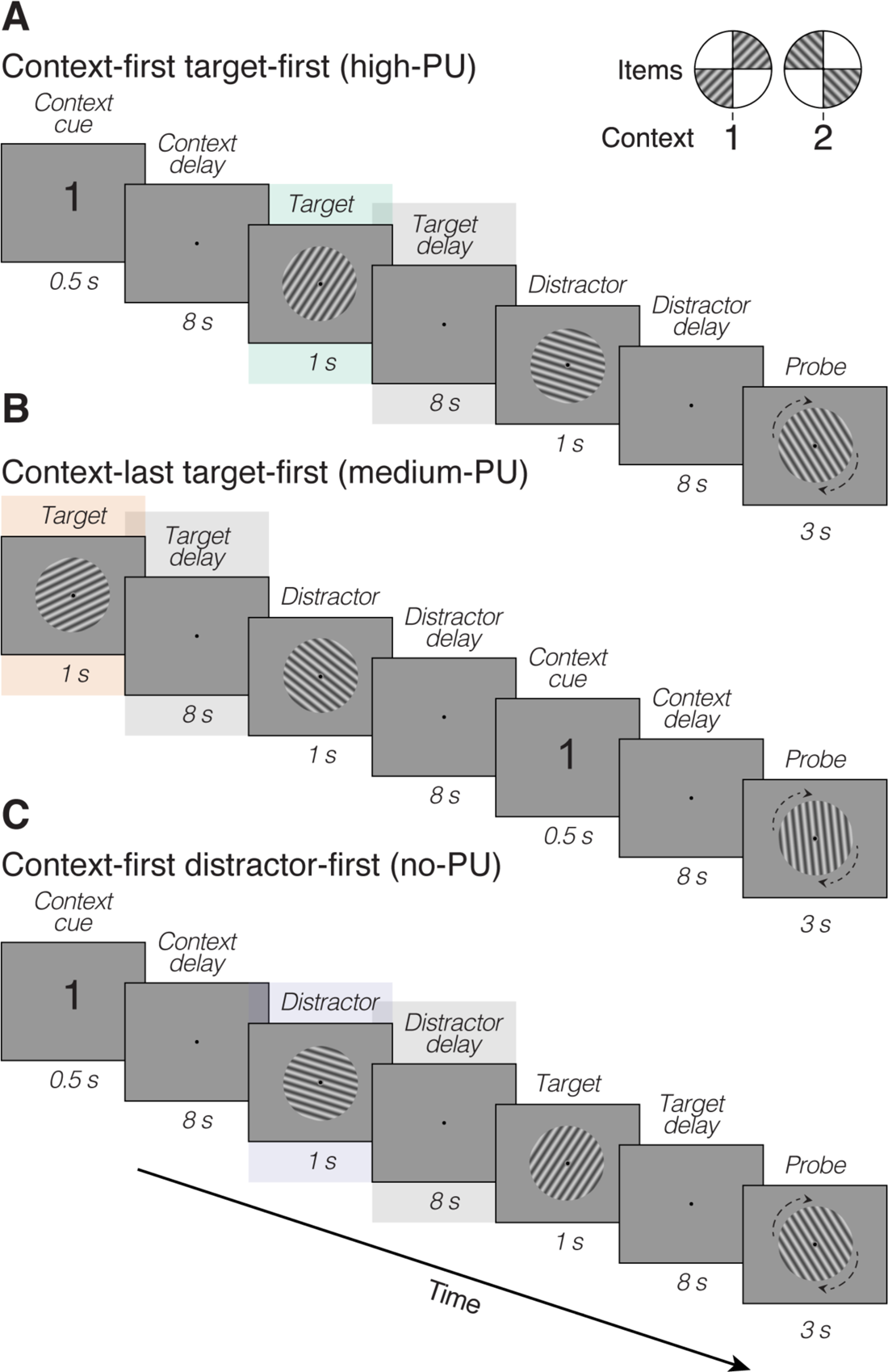
Experimental stimuli. On every trial, participants were presented with two orientation gratings and one context cue, and were asked to match the orientation probe to the remembered target grating. We manipulated predicted utility by varying when the context cue appeared. When the context cue appeared first (context-first trials), participants knew ahead of time which grating to remember (target) and which grating to ignore (distractor). When the context cue appeared last (context-last trials), both gratings had to be remembered until the context cue appeared, at which point the target could be selected. Top-right inset shows schematic indicating the mapping between context cue and orientation category. When context cue is “1”, the target grating is between 0° and 90°. When context cue is “2”, the target grating is between 90° and 180°. ***A,*** context-first trial where the target grating appears first. High predicted utility (high-PU) condition. Green shading indicates which target should be remembered, and gray shading indicates the delay period from which we reconstructed the remembered WM representations. ***B,*** context-last trial where the target grating appears first, although participant is not aware that it is the target. Medium predicted utility (medium-PU) condition. ***C,*** context-first trial where the distractor grating appears first. No predicted utility (no-PU) condition.

In addition to the two orientation gratings, participants were also presented with a digit that acted as a context cue. A single context cue was presented per trial and indicated to participants which orientation was the target. The context cue could either be a 1 or a 2 and this number cued the relevant class of gratings from which the target would come. Specifically, a “1” meant the relevant orientation was a “class-1” grating, corresponding to an arc on the circle falling between 0 and 90 degrees in quadrant 1 and the corresponding degrees between 180 and 270 in quadrant 3 (Fig. 1, top-right inset). If the digit was a “2”, it meant the relevant orientation was a “class-2” grating and would correspond to an arc falling between 90 and 180 degrees in quadrant 2 and the corresponding degrees between 270 and 360 in quadrant 4. To clarify, the context number 1 or 2 did not refer to the order of the appearance of the gratings, as first or second in series, but rather which class of grating was the target. The to-be-remembered orientations never fell within 7° of the vertical cardinal directions or within 17° of the horizontal cardinal direction as this would cause confusion about what context it belonged to.

We manipulated the predicted utility of items by varying the temporal onset of the context cue presentation. The context cue could either appear first, before the orientation gratings, or last, after the orientation gratings. When the context cue appeared first, participants knew ahead of time which orientation grating would be tested (and thus, which grating would be high predicted utility) and which was a distractor. We refer to trials when the context appears first in order by the shorthand “context-first.” When a target item appears in a context-first condition, maintaining this item has high predicted utility because it is 100% likely it will be needed for the response. Conversely, when a distractor appears during a context-first trial, maintaining it has no predicted utility for the upcoming response as there is no chance it will be tested.

When a digit appeared last, participants did not know until the end of the trial which item was to be tested. We refer to this condition as “context-last.” During context-last trials, participants had to encode both orientations into working memory. At presentation, each orientation had the same intermediate predicted utility until the context appeared because each was 50% likely to be useful for the response. Once the context had been presented, participants could select the target item to guide the upcoming response.

In order to relate predicted utility to the fidelity of the remembered angle without the complicating influence of load, we focused on reconstructing the WM representation during the delay period following the first presented grating on each trial. We coded this first delay period as three predicted utility conditions: high predicted utility (high-PU), medium predicted utility (medium-PU), and no predicted utility (no-PU). High-PU delays occurred on *context-first target-first trials* following the presentation of the known target. Medium-PU delays occurred on *context-last target-first trials* following presentation of a grating that would be later cued as the target. No-PU delays occurred on *context-first distractor-first trials* following presentation of a known distractor.

#### Behavioral Procedures

We generated task stimuli in Matlab (The MathWorks) using the Psychophysics Toolbox extensions (Brainard, 1997; Kleiner et al., 2007; Pelli, 1997). All stimuli were centrally presented. In the scanner, stimuli were projected onto a screen at the back of the bore. Stimulus order depended on whether the trial was a context-first or context-last condition. On context-first trials, the context cue (1 or 2) was presented first for 500 ms. An 8-second “context delay” followed the context cue, during which participants fixated a small centrally-located black circle that subtended 0.2° of visual angle.

Next, an oriented sine-wave grating appeared at 25% contrast with a spatial frequency of one cycle per degree. The grating subtended 6.4° of visual angle and flickered at 2 Hz (250 ms on 250 ms off) with the phase of the grating changing 180° on each flicker. This decreased the chances of participants perceiving an afterimage during the delay period. Additionally, there was a 0.15° black fixation circle surrounded by a 0.32° gray ring that participants were instructed to fixate while the grating was on the screen. After the presentation of the first grating, there was another 8-second delay period. Participants were told to only remember the grating if it was the target, indicated by the context cue. After the 8-second delay period, the second orientation grating appeared. This grating was followed by another 8-second delay, in which participants either kept remembering the first grating if that grating had been the target, or remembered the second grating, if the second grating was the target. Participants were instructed to filter the distractor grating out of memory.

Context-last trials were identical to context-first trials except that the orientation gratings appeared first and the context cue appeared last with the context delay preceding the response probe (Fig. 1B).

On both context-first and context-last trials, there was always one class-1 grating and one class-2 grating presented on each trial counterbalanced for order. Class-1 gratings were rendered in 1 of 4 exemplar orientations (20°, 40°, 60°, or 80°) and class-2 gratings were rendered in 1 of 4 different exemplar orientations (100°, 120°, 140°, or 160°). Both gratings were randomly jittered by ± 3° on every trial. The pairing of exemplar orientations within a trial was controlled to be uncorrelated.

At the end of all trials, a response probe appeared for 3 seconds. The response probe had the same sine-wave grating parameters as the class-1 and class-2 gratings, but appeared at an orientation defined as a random offset from the target. Specifically, we randomly selected from a vector of 3°-30° in increments of 3°, and then multiplied that randomly selected orientation by 3 and added it to the target orientation. This procedure ensured that the outcome was a multiple of the target orientation (i.e., that participants would be able to turn the response probe to the actual target orientation). The response probe never appeared at the target orientation. Participants responded by rotating the response probe so that it matched the target grating they were holding in memory. They did this using two keys on an MR-compatible fiber optic response keypad (fORP; Current Designs, Philadelphia, PA).

After a 3-second response period, the response wheel froze and a red feedback line appeared at the target orientation for 440 ms. The feedback line length subtended 3.13° of visual angle. If the participant estimated within 3° of the correct angle, the message, “Correct!” popped up in the middle of the screen instead of the red feedback line. After feedback, a visual noise mask appeared for 60 ms to attenuate any afterimage from the response wheel.

ITIs were sampled randomly from a uniform distribution of 2-6 seconds in increments of 2 seconds, and the same ITI values were used for all participants. Condition order was optimized to first-order counterbalancing to make the design efficient.

In every fMRI session, there were 8 runs of 16 trials each, consisting of 8 context-first trials and 8 context-last trials. In each run, half of the target orientations were class-1 gratings and half were class-2 gratings (i.e., 8 class-1 targets and 8 class-2 targets). Additionally, each exemplar orientation (20°, 40°, 60°, 80°, 100°, 120°, 140°, and 160°) was presented as a target twice in each run, once as a target for context-first trials and once as a target for context-last trials. Each exemplar orientation was also presented as a distractor twice in each run, although this was collapsed across condition type. The frequency that the target orientation appeared first or second in a trial was also counterbalanced within a run. Across runs, orientations of each grating class were fully crossed so that each exemplar orientation from each grating class was presented an equal number of times with each exemplar orientation from the opposite grating class.

### 2.3 fMRI data acquisition

Whole-brain imaging data were acquired at the Brown University MRI Research Facility on a Siemens 3 Tesla MAGNETOM Prisma MRI scanner with a 64-channel head coil. We acquired a high-resolution T1-weighted anatomical image for each subject using a multi-echo MPRAGE (van der Kouwe et al., 2008) pulse sequence (1 mm^3^ resolution; TR = 2530 ms; TE 1.69 ms, 3.55 ms, 5.41 ms, 7.27 ms; sagittal; 176 slices; flip angle = 7°; FoV = 256 mm; TI = 1100 ms). The multi-echo MPRAGE was acquired with AAscout auto-align, which was used to align functional slice placement across sessions. Cortical surface reconstruction was performed on these structural images using FreeSurfer (version 6.0; https://surfer.nmr.mgh.harvard.edu/; Fischl, 2012). Functional images were acquired using a multiband accelerated EPI sequence that allows for simultaneous multi-slice acquisitions of T2*-weighted BOLD images (TR = 2000 ms; TE = 30 ms; flip angle = 86°; slice acceleration factor = 3; 2.4 mm isotropic voxel size; 75 axial slices with 0% interslice skip; matrix size = 86 x 86; FOV = 204 mm). A resting state scan and second T1 MPRAGE were collected in some but not all participants. As these were included at the end of the session and were not analyzed for the present report, we do not detail them further.

Each of the 8 functional runs was 9 minutes and 18 seconds long and consisted of 279 timepoints. We discarded the first 4 volumes of every run to ensure steady-state magnetization. A 4-TR post-cue fixation period was also included at the end of every run.

### 2.4 Preprocessing

We used FreeSurfer FS-FAST software (version 6.0; https://surfer.nmr.mgh.harvard.edu/; Fischl, 2012) to preprocess the fMRI data. We concatenated each subject’s fMRI sessions into one session for the purposes of preprocessing and analysis. Note that session differences were accounted for at the analysis stage. Preprocessing was done in native (individual subject) volume space. We motion corrected each functional run to the first timepoint of the first run, registered each participant’s concatenated functional data to their anatomical, slice-time corrected using FreeSurfer’s default slice time preprocessing method with a multiband factor of 3, and smoothed the functional data at 3 mm full-width half-max.

In order to identify voxels for the generative model, we fit a voxelwise general linear model (GLM) to each participant’s data. Regressors for this model were based on the onsets of stimulus presentation events (presentation of context cues and orientation gratings in both context-first and context-last conditions). Event regressors were modeled as a boxcar of a duration equal to the duration of that event. Regressors were convolved with the SPM canonical hemodynamic response function without derivatives (SPM HRF). Nuisance regressors included a 2nd-order polynomial for detrending and 3 eigenvectors for movement that were derived from the 6 rigid body movement parameters using singular value decomposition.

In order to identify voxels in each participant that responded to stimulus events relative to baseline (i.e., during ITIs), we contrasted stimulus to baseline activation using a t-test at each voxel.

### 2.5 Regions of Interest

We created regions of interest (ROIs) by first masking the task activation map with *a priori* visual areas, taken from a publicly available probabilistic retinotopy atlas (L. Wang et al., 2015). We first projected the Wang parcellation, which is in normalized (fsaverage) surface space, into individual subject surface space. We then created ROIs V1-V3 (separately for left and right hemisphere) out of the Wang parcellation that had been transformed into individual subject space. We chose early visual areas as information has been shown to be encoded in these areas during WM maintenance (Ester et al., 2013; Harrison & Tong, 2009; Lorenc et al., 2018; Pratte & Tong, 2014; Serences et al., 2009; Sprague et al., 2016). In line with the “sensory recruitment hypothesis” (D’Esposito, 2007; Postle, 2006), we specifically chose to use V1-V3 (combining voxels from the dorsal and ventral portions) as these areas have been shown to have high decoding accuracy for orientation representations in WM relative to V3A-V4 (Harrison & Tong, 2009). We did not have *a priori* reason to expect differences among these ROIs. We then transformed each label, which was in individual subject surface space, into volumetric cortical ribbon masks in native space.

Finally, we constructed ROIs by taking the intersection of each *a priori* visual area mask with each participant’s stimulus activity versus baseline significance map using a liberal threshold (p < 0.01, uncorrected). It is important to note that since the size of each ROI depended on each individual participant’s stimuli-vs-baseline activation, the number of voxels in each ROI varied across participants, ranging from 135-434 across all ROIs. In summary, this procedure identified 6 subject-specific V1-V3 ROIs across both hemispheres in each participant.

### 2.6 Mask functional data with Wang intersection ROIs

After preprocessing and ROI creation, we z-scored and detrended the timecourse from each voxel in each ROI. We did this for every participant and every run.

We extracted timepoints for each predicted utility (PU) event: high-PU, medium-PU, and no-PU. For each PU event, we extracted the second to fourth TRs following the offset of the first grating. We chose 2-4 TRs, or 4-8 seconds, after stimulus offset to account for the lag of the hemodynamic response function. Thus, following the logic of prior WM decoding studies (Ester et al., 2015; Harrison & Tong, 2009), though there may be some contribution of the preceding stimulus to the reconstruction, delay activity will be the primary contributor to these reconstructions. These TRs were averaged for each PU event. We created these PU event averages for every participant, for every PU condition, and for every run. We then concatenated the extracted means across all runs.

### 2.7 Statistical analysis

#### Behavioral analyses

To assess memory performance, we fit three established WM models (standard mixture model, swap model, and variable precision model) to each participant’s response error distributions using the MemToolbox (Suchow et al., 2013). The standard mixture model (Zhang & Luck, 2008) assumes participant data falls into one of two types of trials: trials in which participants stored the item in memory, and trials in which the participant did not store the item in memory and guessed. These two types of trials are represented by a mixture of two distributions: a von Mises distribution representing the precision of remembered items, and a uniform distribution representing random guesses. This model results in two parameters: guess rate (*g*) and precision (*sd*). The swap model (Bays et al., 2009) additionally includes a non-target report rate parameter (*b*), representing the proportion of trials in which participants mistakenly remember the distractor item instead of the target item. The variable precision model (Fougnie et al., 2012; van den Berg et al., 2012) assumes that resources are not allocated equally across stored items, nor equivalently across trials. Rather, precision is treated as a random variable drawn from a higher-order distribution with a mean (*sd*_μ_) and standard deviation (*sd_σ_*). We also included a bias parameter in all models so that the central tendency of the fitted data would be able to differ from zero.

We did a model comparison using the corrected Akaike Information Criterion (AIC_c_; Akaike, 1974; Hurvich & Tsai, 1989) to assess which model resulted in the best fit to our data. We determined that the variable precision model performed best for the majority of participants across conditions. To characterize behavioral performance, we used maximum likelihood estimates for the parameters of the variable precision model (*g*, *sd*_μ_, *sd_σ_*, and bias). We fit parameters separately for context-first and context-last trials and separately for each participant. We performed pairwise t-tests between context-first and context-last conditions for each parameter estimate.

#### 2.7.1 Encoding model

We used a Bayesian generative model-based approach introduced by van Bergen et al. (2015) to examine orientation-selective responses during the delay-period. The generative model approach was used because it allows for the decoding of uncertainty (van Bergen et al. 2015). MVPA yields a discrete (relative to continuous) prediction, and the channel response profiles produced by the inverted encoding model (IEM) approach do not necessarily reflect uncertainty. For example, Liu et al. (2018) demonstrated that a change in IEM channel response profile width could result from a change in neural tuning width or simply a change in noise. In the case of zero noise, the channel response profile simply reproduces the channel function assumed by the model. The results produced by the generative model approach, on the other hand, can be interpreted in terms of the certainty by which a neural response indicates the identity of the stimulus stored in WM (Liu et al., 2018; Gardner and Liu, 2019).

The present model assumes voxel responses can be expressed as the linear weighted sum of 8 orientation selective neuronal populations or channels. Each channel was represented by a half-wave rectified sinusoid raised to the 5^th^ power:

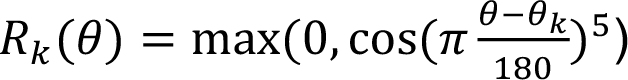

where θ_k_ denotes the preferred orientation of the *k*th population or channel. Channel basis functions were maximally tuned at one of eight equally spaced angles (0°, 22.5°, 45°, 67.5°, 90°, 112.5°, 135°, or 157.5°).

Model fitting and assessment was performed using a 16-fold leave-one-run-out cross-validation procedure, which iteratively partitioned the data into a training set (*B_1_*) and a test set (*B*_2_). The training set (*B_1_*) was expressed as an *n* × *m* matrix, where *n* is the number of trials in the training set and *m* is the number of voxels. Training sets contained 180 trials and were collapsed across PU conditions to avoid introducing any bias. Specifically, training sets included context-first trials where the target appeared first (high-PU condition), context-last trials where the target appeared first (medium-PU condition), and context-first trials where the distractor appeared first (no-PU condition). Although context-last trials where the target appears first are psychologically identical to context-last trials where the distractor appears first (since participants do not know at that point which grating is the target and which is the distractor), we excluded context-last trials where the distractor appears first to keep the number of trials across conditions balanced when training the model. However, we also performed a within-participant replication analysis where we swapped out context-last target-first trials for context-last trials where the distractor came first. This gave us a psychologically identical training set that allowed us to replicate our initial findings using independent within-participant data.

An *n* × *k* channel matrix *C_1_* (where *k* is the number of channels) contained the idealized channel responses for each trial given the presented orientation on that trial (Brouwer & Heeger, 2009; Ester et al., 2013). A *k* × *m* weight matrix *W* related the observed voxel responses to the idealized channel responses:

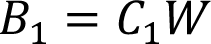

The weight matrix was estimated via ordinary least squares as follows:

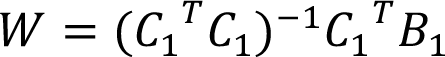

Next, for each trial in the test set we used Bayes’ rule to compute a posterior probability distribution over orientation given the observed pattern of voxel responses.

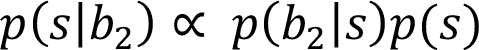

where *s* refers to a particular stimulus value (e.g. 0°) and *b_2_* denotes a single trial/row of the test set BOLD matrix (*B*_2_). We chose the prior *p(s)* to be uniform across angles, since there is no particular reason why *a priori* certain orientations should be favored (van Bergen et al., 2015). To avoid numerical underflow, we performed computations in log space. The log-likelihood of a specific stimulus value given a voxel activation pattern was defined as:

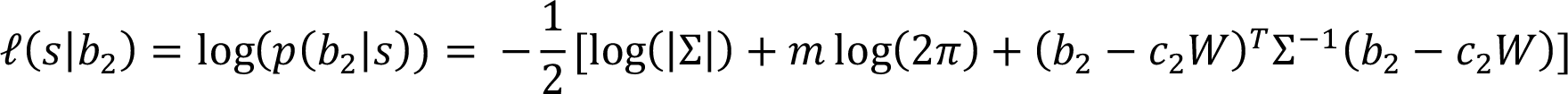

We assume that the bold response follows a multivariate normal distribution with covariance matrix Σ and mean *c_2_W*, where *c_2_* represents a vector of idealized channel responses corresponding with the stimulus value *s* and Σ denotes the *m* × *m* voxel covariance matrix estimated from the training set.

As computing the log-likelihood requires computing the inverse of the voxel covariance matrix, we performed a procedure described in Naselaris et al. (2009) if Σ was ill-conditioned. First, we performed principal component analysis on the predicted responses (B^^^_1_) for the training set. *p* components were chosen with the constraint that each component explained at least 5% of the total variance. Using the resulting *m* × *p* projection matrix, we then projected the predicted (B^^^_1_) and actual (*B_1_*) responses from the training set onto the first *p* principal components. These dimensionality-reduced responses were then normalized to unit length:

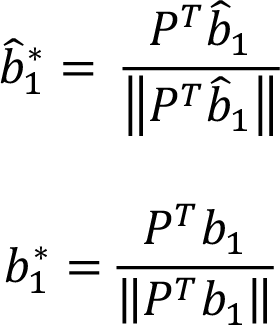

where *P* represents the computed projection matrix, b^^^_1_ and b_1_ respectively denote a single row from the predicted (B^^^_1_) and actual (*B_1_*) response matrix, and b^^^_1_^∗^ and b_1_^∗^ respectively denote a single row from the dimensionality-reduced *n* × *p* predicted (B^^^_1_^∗^) and actual (B_1_^∗^) response matrix. The voxel covariance matrix was then computed as:

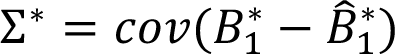

where Σ^*^ is a *p* × *p* matrix.

During the testing phase, the same procedure (projection onto the first *p* components using the projection matrix obtained from the training set followed by normalization) was performed on the predicted and actual responses for the test set. Thus, the log-likelihood of a specific stimulus value given the dimensionality-reduced pattern of voxel responses was re-expressed as:

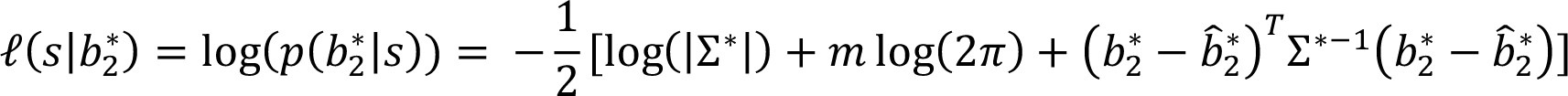

To assess whether an area contains stimulus-specific information across trials, we then circularly shifted trial-level log-likelihoods so that the center value corresponded with the presented stimulus on that trial. Probability densities, as normalized ratios, must be averaged using the geometric mean rather than the arithmetic mean (Fleming & Wallace, 1986; Nelson, 2017). This is equivalent to computing the arithmetic mean of the log-likelihood and then exponentiating the result. Consequently, we averaged the shifted log-likelihoods across trials to produce a subject-level log-likelihood. We then averaged subject-level log-likelihoods to form a group-level log-likelihood, which was then exponentiated and normalized to compute a posterior probability distribution.

To test that our probability density distributions were significantly different than noise, we performed permutation testing. We implemented a data-wise permutation scheme (Etzel & Braver, 2013) that permuted the orientation labels within a cross-validation fold prior to model training and assessment. We performed 1000 permutations. Each of these permutations followed the same procedure as described above: 1. encoding model training and testing performed separately for each subject, ROI, and hemisphere; 2. log-likelihoods shifted and averaged across trials and subjects and then exponentiated and normalized to compute a geometric mean probability distribution. We then estimated the significance of reconstructed orientations by comparing the probability of the presented stimulus (corresponding with p(0° | *b*)) for circularly shifted distributions) to our empirically estimated null distribution (Golland & Fischl, 2003). A p-value was computed as [(# of permutation probabilities ≥ actual probability) + 1] / (N permutations + 1) (Phipson & Smyth, 2010). We performed this procedure for every ROI, hemisphere, and predicted utility condition.

In addition to performing permutation tests for our main TR window of interest (2-4 TRs, or 4-8 seconds after delay onset of the first grating), we also performed permutation testing for three additional subsequent TR windows (TRs 3-5 (6-10 seconds), 4-6 (8-12 seconds), and 5-7 (10-14 seconds)), in order to discern distractor decodability in the no-PU condition.

To compare WM representations across predicted utility conditions we examined three measures of interest: two complementary measures of decoding uncertainty (entropy and posterior standard deviation), both of which served as our measures of stimulus representation fidelity, and posterior probability of the presented/remembered stimulus given the observed BOLD activity.

For our first measure of stimulus uncertainty/fidelity, we computed the entropy of the subject-level posterior distributions, computed via geometric averages as described above. We then computed the entropy of these subject probability distributions with the following formula:

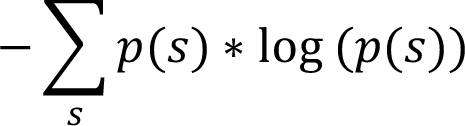

where *s* represents a particular orientation and *p(s)* denotes the probability of that orientation.

Entropy accounts for any small change over the whole posterior and is computed without particular reference to a mean angle, making it a less sensitive measure for variance around the presented angle. Thus, we also computed the circular posterior standard deviation (posterior SD), which has been used as an estimate for stimulus uncertainty in prior work (van Bergen et al., 2015).

We tested whether entropy and posterior SD differences exist across predicted utility conditions (separately) by fitting linear mixed effects models using the lmerTest package (version 3.1-2; Kuznetsova et al., 2017) in RStudio 1.2.5001 (R Core Team, 2018). For each measure, we implemented two models to test for significant differences between conditions. Model 1 included main effects of predicted utility condition, ROI, and hemisphere, with ROI and hemisphere meant to be nuisance regressors. Subject was entered as a random effect to account for between-subject variance. Model 2 included main effects of predicted utility condition and ROI, as well as their interaction. The lmerTest package calculated denominator degrees of freedom and p-values for each model via Satterthwaite’s degrees of freedom method. Each linear mixed effects model computed an average for every subject and incorporated between-subject variance in the model.

In addition to computing entropy and posterior SD across trials, we also computed these measures of uncertainty (entropy and posterior SD) at a single trial level. This allowed us to differentiate between two factors that could potentially influence uncertainty: (1) if single-trial posteriors have large uncertainty (i.e., large entropy or large posterior SD), then the average of those posteriors will also have large uncertainty, or (2) if single-trial posteriors have low uncertainty but large variance in the mean (i.e., in the estimate of the predicted orientation), then the averaged posteriors will also reflect large uncertainty.

Entropy and posterior SD were computed in the same way as described above, except that each was computed from an individual trial-level posterior distribution, instead of from a trial-averaged posterior distribution. We tested whether there were differences in trial-level entropy and posterior SD (separately) across predicted utility conditions by implementing the same two linear mixed effects models as described for the trial-averaged posteriors above.

The posterior probability of the presented/remembered stimulus can be thought of as the probability of the true stimulus given the observed BOLD activity. To calculate this, we took the shifted and averaged posterior distribution for each subject and then extracted the probability at the center value of 0°. A higher value is indicative of greater information about the stimulus. We tested for significant differences in posterior probability of the presented/remembered stimulus between predicted utility conditions by performing the same two linear-mixed models as described for entropy.

Finally, although our task design allows us to look at how predicted utility modulates fidelity independent of load, there are differences across condition in cue order. Specifically, high-PU and no-PU trials had a pre-cue, while medium-PU trials did not. Thus, to differentiate between plausible mechanisms that might contribute to predicted utility modulating fidelity, we also performed a time window analysis, with the idea that attentional processes may affect fidelity earlier in the delay window, while gating may affect fidelity later in the delay. We specifically looked at TR windows later in the delay period after the first grating had been presented (TRs 3-5 (6-10 seconds), 4-6 (8-12 seconds), and 5-7 (10-14 seconds)) and examined whether entropy, posterior SD, and the posterior probability of the presented/remembered stimulus differed between each TR window.

## 3. Results

### 3.1 Behavioral Results

Behavioral performance indicated that participants performed the task accurately and took advantage of contextual information when it is was presented first (i.e., context-first trials). We compared response accuracy and precision between context-first and context-last conditions using estimates from the variable precision model with bias (Fougnie et al., 2012; van den Berg et al., 2012), which had the best fit to our data (see Methods). This model provides an estimate of precision in terms of the standard deviation of the participant’s error distribution on the memory task, as well as an estimate of the standard deviation of the higher-order variability distribution. That is, since precision is variable and not fixed, precision is considered to itself be a random variable that is drawn from a distribution with a mean (mean precision, *sd*_μ_) and a standard deviation (*sd_σ_*). Our precision results were comparable to other studies that also fit continuous report data using the variable precision model (Fougnie et al., 2012; van den Berg et al., 2012). Memory was more precise in the context-first condition relative to the context-last condition, and had less variability in the higher-order variability distribution (see *sd*_μ_ and *sd_σ_* in Table 1).

**Table 1.** Variable precision model behavioral results for context-first and context-last trials, as well as target-first/target-last results for each trial type. *sd*μ is mean precision, *sdσ* is precision variability, and *g* is guess rate.

| | $sd_{\mu} (^{\circ})$ | $sd_{\sigma} (^{\circ})$ | $g$ | Bias ($^{\circ}$) |
| --- | --- | --- | --- | --- |
| Context-first | 7.236 | 2.533 | 0.02 | 0.368 |
| Target-first | 8.121 | 2.681 | 0.02 | 0.4311 |
| Target-last | 5.906 | 2.393 | 0.02 | -0.086 |
| Context-last | 8.817 | 4.01 | 0.002 | 0.382 |
| Target-first | 10.331 | 2.917 | 0 | -0.063 |
| Target-last | 6.258 | 4.597 | 0.004 | 0.302 |

A paired t-test showed that *sd*_μ_ was significantly different between context-first and context-last: *t*_(6)_ = −3.231, *p* < 0.05, Cohen’s d = −1.221, although *sd_σ_* was not significantly different (*t*_(6)_ = −0.899, *p* = 0.40, Cohen’s d = −0.34). Note that the guess rate (*g*) was very low overall (0.02 for context-first and 0.002 for context-last) and did not differ significantly between the context-first and context-last conditions (guess rate: *t*_(6)_ = 1.982, *p* = 0.095, 0.749). Bias was also low overall (< 0.5°) and did not differ between context-first and context-last trials (bias: *t*_(6)_ = − 0.022, *p* = 0.98, −0.009).

Because time is known to tax WM (Park et al., 2017; Zhang & Luck, 2009), we tested whether there were precision differences when the target appeared first versus last. When collapsing across context-first and context-last trials, there was a trending effect of order where *sd*_μ_ was greater for targets that appeared first compared to targets that appeared last (*t*_(21)_ = −1.957, *p* = 0.0637, Cohen’s d = 0.906). However, there was no interaction between trial type (context-first vs. context-last) and target position (*t*_(21)_ = −1.16, *p* = 0.259).

The condition difference in WM precision indicates that participants took advantage when the context appeared first in order to control memory. However, it is not possible to attribute this difference in precision to predicted utility of the items independent of load, as WM load differs between context-first (one item) and context-last (two items), when participants use the context proactively. Thus, we turned to fMRI to assess how the fidelity of WM representations in the first item delay were influenced by predicted utility when load was controlled.

### 3.2 Stimulus encoding model

To investigate the effect of predicted utility on the fidelity of WM representations in the brain, we focused analyses on the delay period following presentation of the first oriented grating stimulus. In this way, the load was comparable between the critical high- and medium-PU conditions. We fit an orientation encoding model to our fMRI data that uses information about voxel orientation preferences to generate a probability distribution over orientation for each trial given the observed pattern of activity (see Methods). We generated a group-level probability distribution for each condition (high-PU, medium-PU, and no-PU) and ROI (V1, V2, V3 in left and right hemisphere). If an ROI contains orientation-selective information about a stimulus encoded into WM, then this shifted and averaged distribution should peak at or near 0°.

Across all ROIs, the probability of the presented stimulus given the observed activity (p(0° | b) for shifted distributions) was highest for the high-PU condition (Fig. 2), followed by medium-PU, and lowest for no-PU. Non-parametric permutation tests revealed a robust representation of the orientation stored in WM in the high-PU and medium-PU conditions for all ROIs (Table 2). The no-PU condition was weakest, as expected, but a trace reconstruction above chance was found in left hemisphere V2 and V3 and right hemisphere V1 and V2. This might reflect stimulus-related activity stemming from the sluggish hemodynamic response.

**Figure 2.**
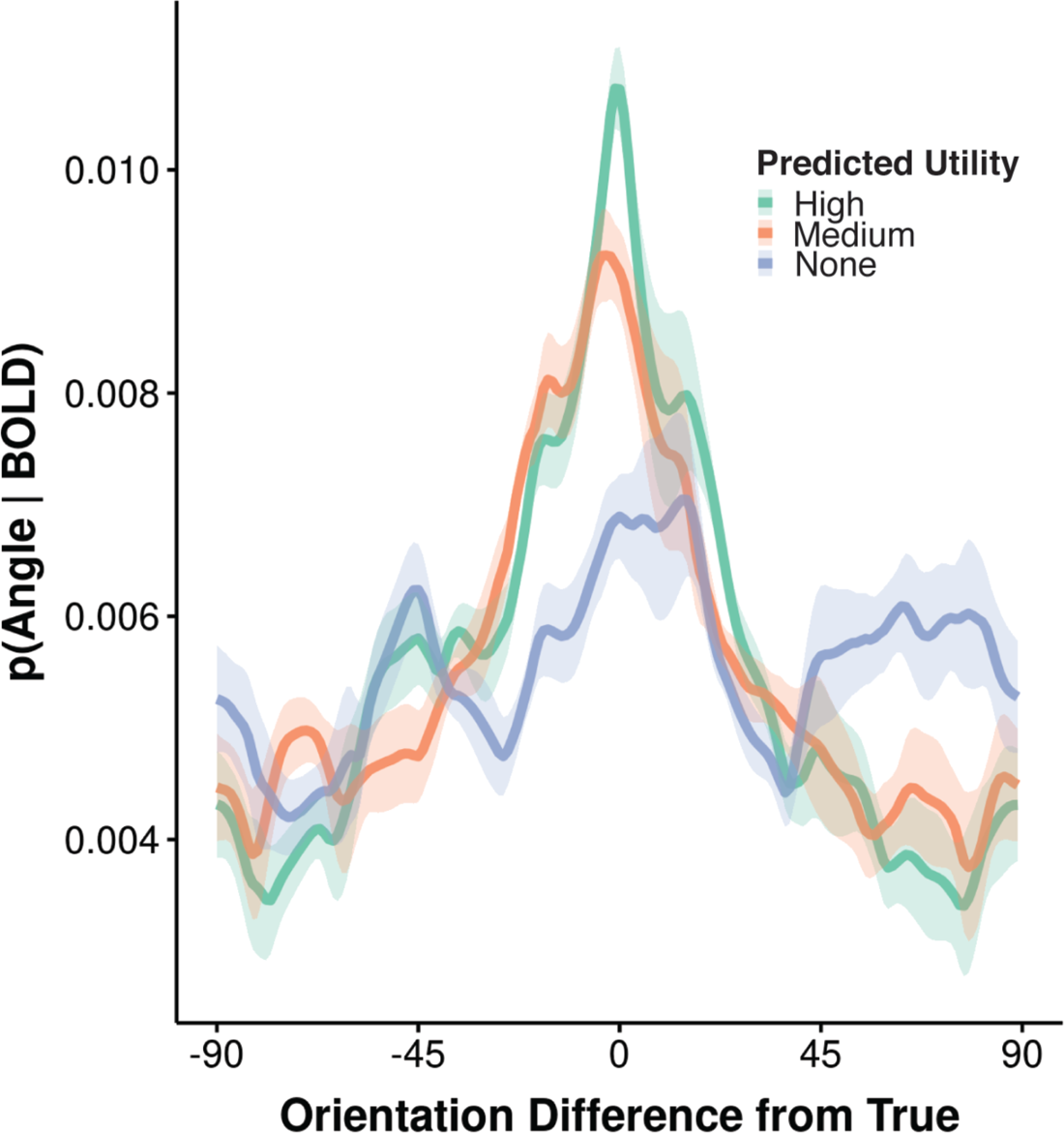
Posterior reconstruction results. Posterior probability distributions over orientation that have been circularly shifted so that 0° corresponds with the actual stimulus presented and averaged across subjects. These group-level posteriors indicate which orientation is most probable given the observed pattern of activity in bilateral V1-V3 during the delay period. Here, we observe a parametric effect of predicted utility on stimulus representation strength, where the posterior probability of the presented/remembered stimulus is highest for high predicted utility, followed by medium predicted utility, and lowest for no predicted utility. Shaded ribbon indicates a bootstrap estimate of SEM.

**Table 2.** P-values from non-parametric permutation tests comparing probability of true posteriors to null distribution for each predicted utility condition. All p-values were corrected for multiple comparisons using false discovery rate (FDR) (Benjamini and Yekutieli, 2001).

| ROIs | Predicted Utility |  |  |
| --- | --- | --- | --- |
|  | High | Medium | None |
| V1 |  |  |  |
| Left hemisphere | 0.0016 | 0.0016 | 0.0529 |
| Right hemisphere | 0.0016 | 0.0016 | 0.0124 |
| V2 |  |  |  |
| Left hemisphere | 0.0016 | 0.0016 | 0.0064 |
| Right hemisphere | 0.0016 | 0.0016 | 0.0124 |
| V3 |  |  |  |
| Left hemisphere | 0.0016 | 0.0016 | 0.0041 |
| Right hemisphere | 0.0016 | 0.0041 | 0.4915 |

We confirmed this by performing permutation tests on TRs that were shifted later (i.e., TRs 3-5, 4-6, and 5-7, in addition to the 2-4 TR window of the current analysis), to see if the distractor could still be decoded at these later timepoints. The distractor was decodable in all but one of the ROIs at the first time window (TRs 3-5), but not decodable in any of the ROIs by the last time window (TR 5-7). This suggests that the earlier decodability could have been due to leakage from the stimulus. Overall, however, we find that fidelity of the maintained representation within visual cortex to be related to predicted utility.

To test our hypothesis that predicted utility increases fidelity of WM representations (i.e., reduces uncertainty), we computed the entropy and standard deviation of posterior distributions (posterior SD), which provide complementary measures of uncertainty based on the evidence for all the possible orientations. Higher posterior SD indicates greater width, or greater uncertainty, in an error distribution around the peak estimated probability. Higher entropy can be interpreted in a way similar to higher posterior SD of an error distribution, except that it does not rely on the assumption of any particular parametric shape (e.g., Gaussian). Thus, higher entropy also corresponds with greater uncertainty in the neural representation of remembered stimuli.

We observed a parametric pattern of results where, when collapsed across ROIs and hemisphere, entropy was lowest (i.e., lowest uncertainty) for high-PU, followed by medium-PU, and highest for no-PU (mean = 5.09, 5.11, 5.13, respectively) (Fig. 3A). This pattern was the same for all ROIs and hemispheres. We validated these observations by fitting a linear mixed effects model using the lmerTest package (Kuznetsova et al., 2017) in RStudio 1.2.5001 (R Core Team, 2018). We included PU condition, ROI, and hemisphere as main effects. Subject was included as a random effect. All comparisons were FDR corrected unless otherwise stated. This model revealed that entropy for high-PU was lower than both medium-PU (*t*(119) = 2.25, P = 0.026) and no-PU (*t*(119) = 5.2., P = 2.52 x 10^-6^). Likewise, medium-PU entropy was significantly lower than no-PU entropy (*t*(119) = 2.95, P = 0.006). There was no interaction between PU condition and ROI (F(4, 119) = 0.504, p = 0.733; uncorrected). Thus, as we predicted, the fidelity of items in memory was modulated by their predicted utility.

**Figure 3.**
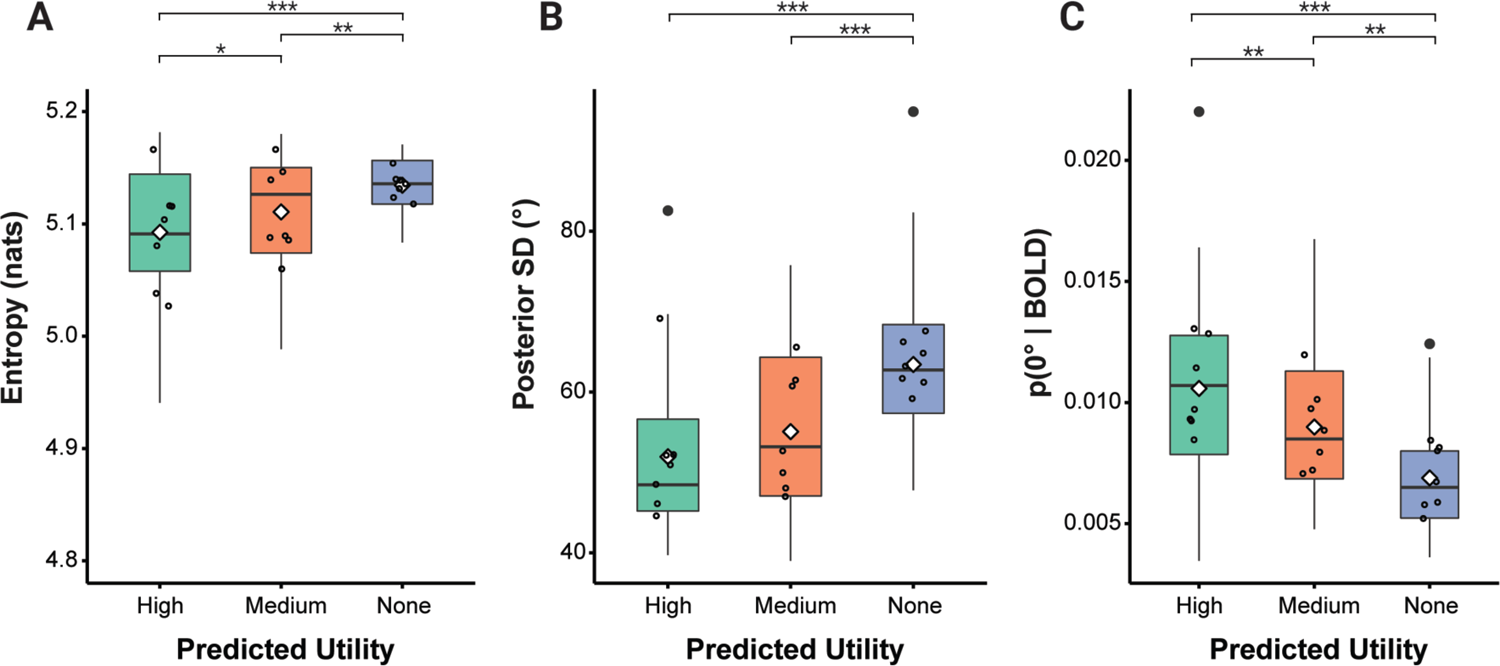
Predicted utility influences neural uncertainty and posterior probability of the stimulus. Each boxplot shows the distribution (interquartile range) of entropy, posterior SD, or p(0°|BOLD) values across participants for a single predicted utility condition. The white diamond indicates the mean, the middle black line indicates the median, top and bottom of box indicate upper and lower quartile, and whiskers indicate the full range of values excluding outliers. Larger filled black circles indicate outliers. Smaller unfilled black circles are individual datapoints. Values have been collapsed across hemisphere and ROI (V1-V3). ***A***, stimulus uncertainty (entropy) results of subject-level posterior probability distributions. ***B***, stimulus uncertainty (posterior SD) activity results. ***C***, posterior probability of the presented stimulus given the observed BOLD activity results. The probability of each subject-level shifted posterior was extracted at the center value (0°). A higher probability value at 0° indicates greater information about the presented orientation and thus greater representation strength. Asterisks denote significance (*p < 0.05, **p < 0.01, ***p < 0.001; FDR corrected).

We observed a similar parametric pattern of results for posterior SD as we did for entropy. When collapsed across ROIs and hemisphere, posterior SD was lowest (i.e., lowest uncertainty) for high-PU, followed by medium-PU, and highest for no-PU (Fig. 3B). This pattern was the same for all individual ROIs and hemispheres except for the right hemisphere of V1, in which medium-PU had the lowest posterior SD instead of high-PU. We validated these findings by fitting the same linear mixed effects model used to fit entropy. High-PU posterior SD trended significant relative to medium-PU (*t*(119) = 1.90, P = 0.06), and was significantly lower than no-PU (*t*(119) = 6.98, P = 5.41 x 10^-10^). Additionally, medium-PU was significantly lower than no-PU (*t*(119) = 5.08, P = 2.11 x 10^-6^). Thus, overall these results reflect the same general pattern as observed with entropy.

The preceding analysis found that decoded entropy and posterior SD are parametrically modulated by predicted utility. We next sought to assess the relationship between predicted utility and the posterior probability of the presented/remembered stimulus given the pattern of BOLD activity. A higher probability indicates that the voxel activity carries more information about the presented/remembered stimulus orientation. Overall, we observed a similar pattern of results as we did for entropy. The posterior probability of the presented stimulus was highest for high-PU, followed by medium-PU, and lowest for no-PU (mean = 0.011, 0.009, 0.007) (Fig. 3C). This was true for all ROIs for both left and right hemisphere.

We confirmed these results with a linear-mixed model that was identical to the models used to fit entropy and posterior SD. This model demonstrated that the probability of the presented/remembered stimulus for high-PU was significantly different than both medium-PU (*t*(119) = −2.63., P = 0.01) and no-PU (*t*(119) = −6.11., P = 3.84 x 10^-8^). This model also revealed a significant difference between medium-PU probability and no-PU probability (*t*(119) = −3.48, P = 0.001). There was no interaction between PU condition and ROI (F(4, 119) = 0.611, p = 0.656; uncorrected). Thus, memory for items encoded with higher predicted utility carry progressively more information about those items in the brain.

### 3.3 Within-participant replication analysis

As a complementary analysis, we also tested the distractor in context-last trials (Fig. 4). For this analysis, our training set included the same context-first target-first and context-first distractor-first trials as before, but we now included context-last trials where the distractor came first, and left out context-last trials where the target came first. From the participant’s perspective, context-last trials where the distractor comes first are not psychologically different than trials where the target comes first, allowing us to replicate our findings within-participant on independent trials of the medium-PU condition. We did not include these context-last distractor-first trials in the original analysis to ensure that the number of trials from each condition type was balanced during model training (see Methods).

**Figure 4.**
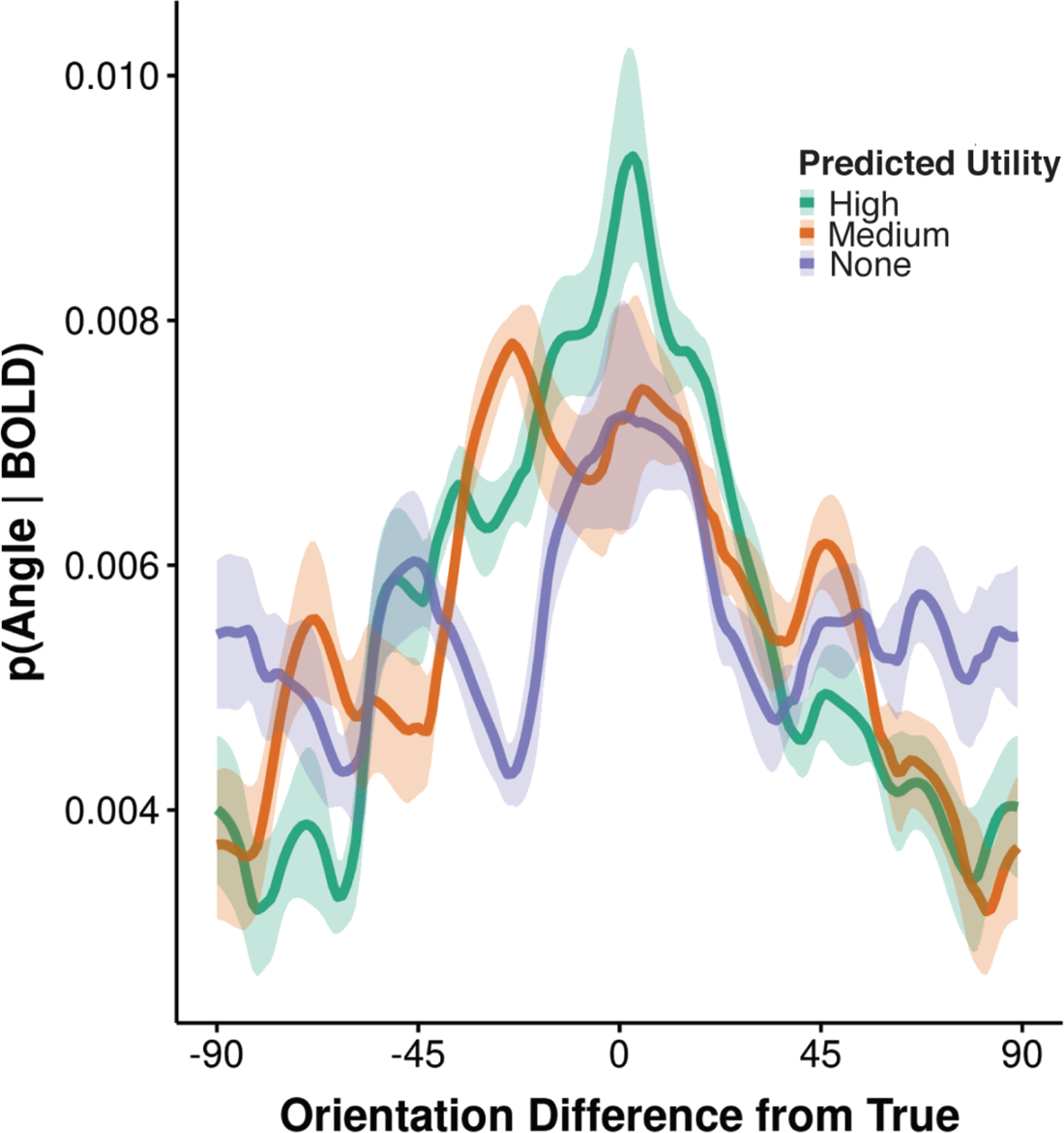
Replication analysis results. Group-level shifted and averaged posterior probability distributions for the replication analysis. Here, we observe that the posterior of the presented/remembered stimulus is highest for high predicted utility, but does not appear to differ between medium predicted utility and no predicted utility. Shaded ribbon indicates a bootstrap estimate SEM.

Non-parametric permutation tests revealed a robust representation of the orientation stored in WM in the high-PU and medium-PU conditions for all ROIs, as we observed in the main analysis, but also in the no-PU condition for all ROIs, which we only observed for 4 out of 6 ROIs in the main analysis (Table 3). As we discussed above, this effect on no-PU trials may be due to vestigial stimulus-related activity.

**Table 3.**
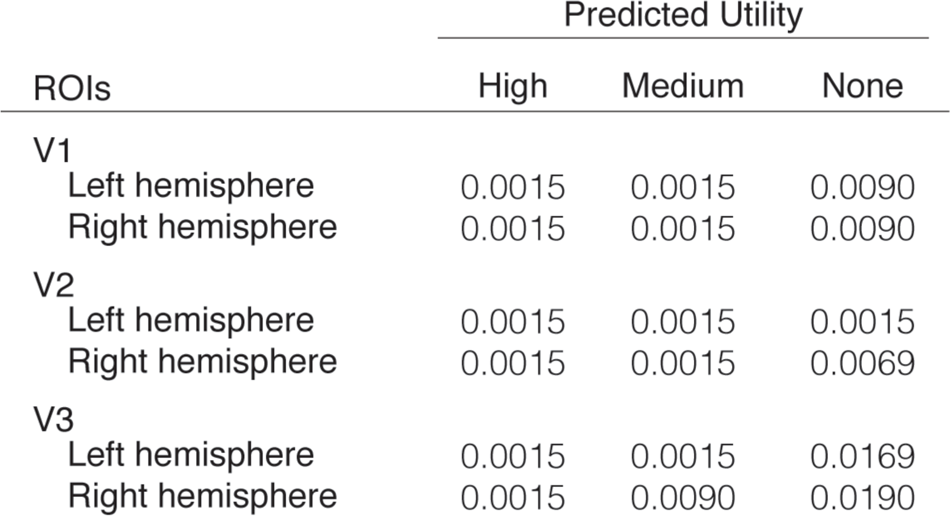
Permutation test results from replication analysis. P-values come from non-parametric permutation tests comparing probability of true posteriors to a null distribution for each predicted utility condition. P-values have been corrected using false discovery rate (FDR).

| ROIs | Predicted Utility |  |  |
| --- | --- | --- | --- |
|  | High | Medium | None |
| V1 |  |  |  |
| Left hemisphere | 0.0015 | 0.0015 | 0.0090 |
| Right hemisphere | 0.0015 | 0.0015 | 0.0090 |
| V2 |  |  |  |
| Left hemisphere | 0.0015 | 0.0015 | 0.0015 |
| Right hemisphere | 0.0015 | 0.0015 | 0.0069 |
| V3 |  |  |  |
| Left hemisphere | 0.0015 | 0.0015 | 0.0169 |
| Right hemisphere | 0.0015 | 0.0090 | 0.0190 |

In terms of stimulus uncertainty and probability of the presented/remembered stimulus given the observed activity, we reproduced the pattern of findings from the above analyses with the exception that the probability for the presented/remembered stimulus for medium-PU was not reliably different than no-PU. High-PU entropy was significantly lower than both medium-PU *(t*(119) = 3.34, P = 0.002) and no-PU *(t*(119) = 5.5, P = 6.65 x 10^-7^), and medium-PU entropy was significantly lower than no-PU entropy *(t*(119) = 2.16, P = 0.033) (Fig. 5A). High-PU posterior SD was significantly lower than medium-PU *(t*(119) = 3.68, P = 0.0005) and no-PU *(t*(119) = 5.87, P = 1.2 x 10^-7^), and medium-PU was significantly lower than no-PU *(t*(119) = 2.19, P = 0.03). The high-PU probability of the presented/remembered stimulus was significantly higher than both medium-PU *(t*(119) = −3.84, P =0.0005) and no-PU *(t*(119) = −3.69, P = 0.0005) (Fig. 5B). However, we observed no significant differences between the probability of the presented/remembered stimulus for medium-PU compared to no-PU *(t*(119) = 0.15, P = 0.884), as we had in the original analysis. Thus, overall, this replication analysis confirms that the fidelity of stimulus probability distributions encoded by visual cortical activity is influenced by predicted utility.

**Figure 5.**
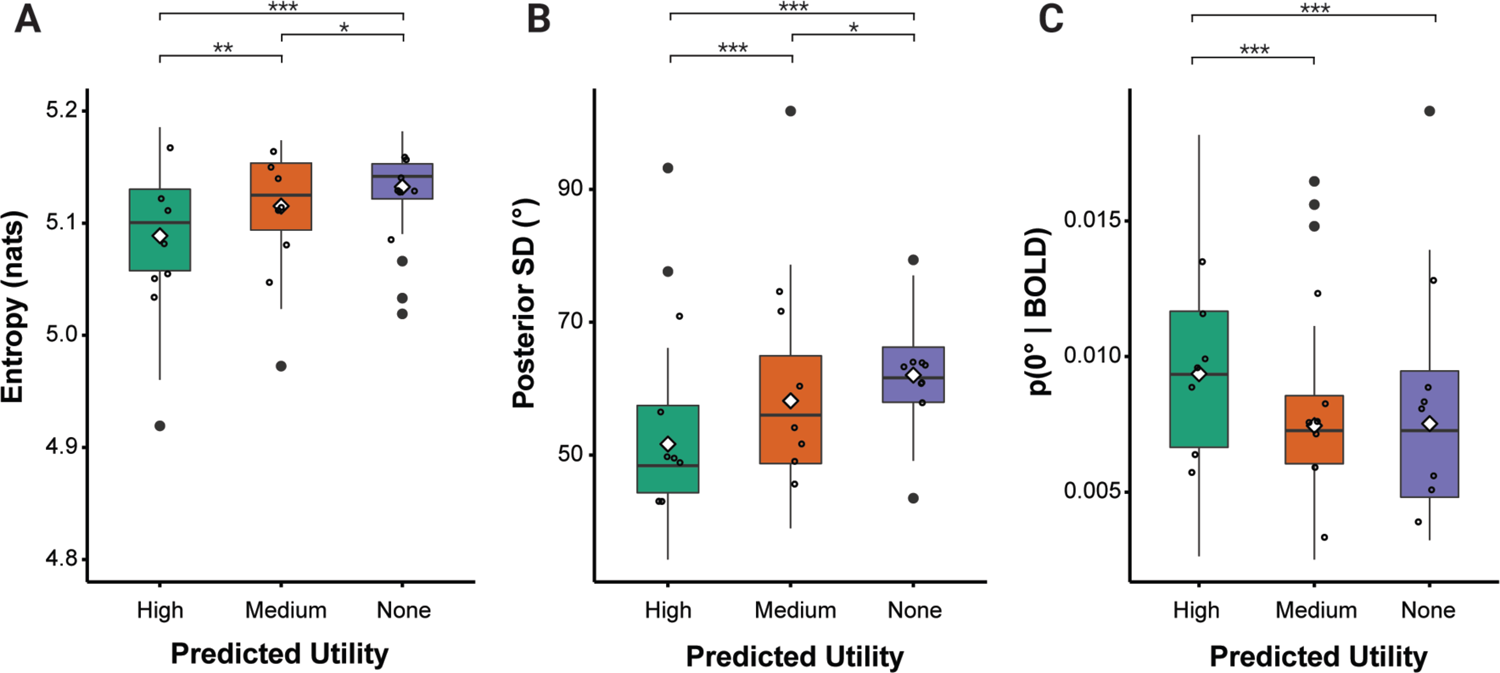
Replication analysis results. ***A***, stimulus uncertainty (entropy) of subject-level posterior probability distributions for high, medium, and no predicted utility. ***B,*** stimulus uncertainty (posterior SD) results. ***C,*** posterior probability of the presented stimulus given the observed BOLD activity results. The probability of each subject-level shifted posterior probability distribution was extracted at the center value (0°) for high, medium, and no predicted utility. Boxplots show mean (white diamond), median (middle black line), quartiles (boxes), range (whiskers), and outliers (black circles). Values have been collapsed across hemisphere and ROI (V1-V3). Asterisks denote significance (*p < 0.05, **p < 0.01, ***p < 0.001; FDR corrected).

### 3.4 Trial-level uncertainty analysis

As discussed in the Methods, we also performed trial-level analyses for both uncertainty measures (entropy and posterior SD) in order to differentiate whether the degree of uncertainty observed across trials was due to the same degree of uncertainty trial-to-trial or to high variance in the mean of the posterior. We performed the same linear mixed effects models that were used in the trial-averaged measures. All p-values have been FDR-corrected.

We initially tested entropy. High-PU was still significantly lower than medium-PU (*t*(8057) = 4.74, P = 6.39 x 10^-6^), but was not significantly different than no-PU (*t*(8057) = 1.28, P = 0.20). Additionally, medium-PU had greater entropy than no-PU (*t*(8057) = −3.47, P = 0.0008). However, as explained in the Methods, entropy is computed without reference to a mean angle, and thus accounts for any small change over the entire posterior. This means that in distributions with multiple peaks, entropy will be the same regardless of how far apart the peaks are from each other. This is an issue because we are interested in how uncertainty varies about a mean angle: peaks that are farther from the presented angle should indicate greater uncertainty than peaks closer to the presented angle. That is, more distance between peaks should correspond with greater uncertainty. Using entropy for trial-averaged posteriors did not pose as much of an issue because most distributions are unimodal. However, the majority of single-trial level distributions are multimodal. As a consequence, at the single-trial level, entropy may not provide a sensitive test of uncertainty.

Thus, we complemented entropy with an analysis of posterior SD for trial-wise analyses. Posterior SD is computed based on the variance in angles about the mean direction and so represents the uncertainty around the presented angle. Thus, we also used posterior SD as a more sensitive test of whether there are wider posteriors at the single-trial level. As expected, this analysis mirrors that found in the trial-averaged analysis. In particular, high-PU had significantly lower posterior SD than medium-PU (*t*(8057) = 3.52, P = 0.0007) and no-PU (*t*(8057) = 4.61, P = 1.24 x 10^-5^). Additionally, although medium-PU had lower overall standard deviation relative to no-PU, this difference was not significant (*t*(8057) = 1.09, P = 0.28, uncorrected). Thus, these results suggest that the trial-level effects are likely due to changes in uncertainty on every trial due to predicted utility.

We also performed trial-level analyses for our uncertainty measures (entropy and posterior SD) for our within-subject replication analysis. We focus on the posterior SD results for the reasons noted above. For trial-level posterior SD, we did not observe a significant difference between high-PU and medium-PU *(t*(8057) = −1.63, P = 0.10), but did observe that high-PU posterior SD was significantly lower than no-PU *(t*(8057) = 2.35, P = 0.03). We also observed that medium-PU posterior SD was significantly lower than no-PU posterior SD *(t*(8057) = 3.98, P = 0.0002). Thus, overall, our trial-level posterior SD results suggest that visual cortical activity is influenced by predicted utility, although perhaps not to the full parametric effect that was observed from our trial-averaged results.

### 3.5 Time window analysis

To examine what mechanisms might be contributing to predicted utility’s effect on fidelity, we performed a time window analysis looking at TRs later into the first-grating delay period. In particular, we reasoned that an attentional pre-cueing effect would be evident early in the WM delay, even at the point of stimulus processing. By contrast, the benefits of gating could emerge later, particularly when noise has greater opportunity to disrupt maintenance of items that are not gated. Our original analysis only looked at a delay window that was predicted to be the peak of the hemodynamic response from 4-8 seconds after delay onset. So, we performed additional analyses looking at sequential temporal windows later in the delay (TRs 3-5 (6-10 seconds), 4-6 (8-12 seconds), and 5-7 (10-14 seconds)).

We found that the posterior probability of the presented/remembered stimulus (i.e., p(0°|BOLD)) was differentiated by utility in a parametric way only at the latest TR 5-7 window (see Fig. 6C). This TR 5-7 window was 10-14 seconds after the onset of the first delay, and 1-5 seconds after the second grating was presented, meaning there might have been interference from the second grating in this interval. This suggests that the strength of the representation of the first-presented target grating is modulated by predicted utility later in the delay when other information has an opportunity to disrupt it, as would be predicted by a WM gating account.

**Figure 6.**
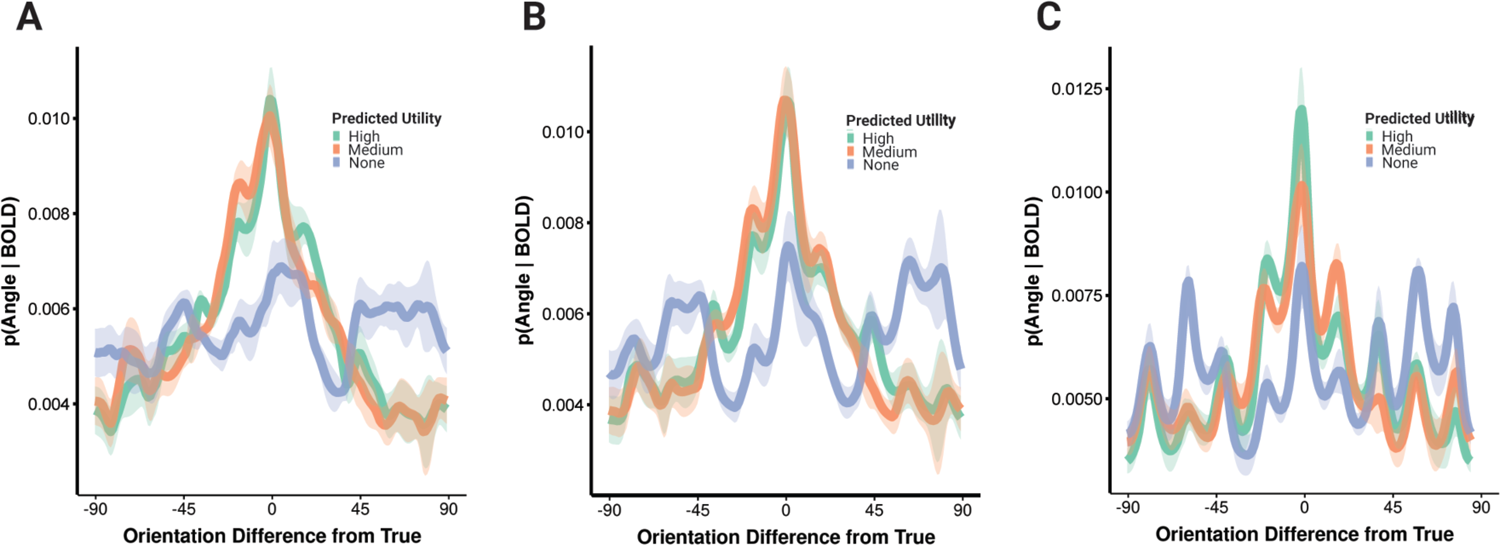
TR window analysis results. Group-level shifted and averaged posterior probability distributions for ***A***, TRs 3-5 (6-10 seconds), ***B,*** TRs 4-6 (8-12 seconds), and ***C,*** TRs 5-7 (10-14 seconds) after the delay onset of the first grating presentation. Shaded ribbon indicates a bootstrap estimate SEM.

To verify whether there were differences in how predicted utility level modulated representation strength depending on the TR window, we performed a linear mixed effects model testing the interaction between PU level and TR window on the probability of the presented/remembered orientation, or p(0°|BOLD) (note: we did not correct for multiple comparisons because the TR windows are not independent of each other). When high PU and the TR 3-5 window were used as reference levels, we observed a trending level of significance where the difference between going from the 3-5 TR window to the 5-7 TR window resulted in a greater posterior probability (p(0|BOLD)) for high-compared to medium-PU *(t*(371) = −1.919, P = 0.0557). Further, when we used TRs 4-6 as a reference level, we observed a significant effect where the difference between TRs 4-6 and 5-7 increased the posterior probability for high-versus medium-PU *(t*(371) = −2.194, P = 0.0288). We also observed a significant interaction between TRs 4-6 and 5-7 for high-versus medium-PU for posterior SD *(t*(371) = 2.499, P = 0.0129), although not for entropy.

## 4. Discussion

In this experiment, we sought to test the hypothesis that the brain can control the fidelity of visual information held in working memory based on that item’s predicted utility for subsequent behavior. Importantly, though prior work has shown that memory is improved for higher-priority items and fidelity is likewise better, it has not been established whether there are mechanisms that can enhance fidelity for an item in a graded fashion based on the predicted utility of maintaining that item itself, as opposed to the utility of maintaining it relative to other concurrently maintained items. This is important to resolve because the answer to this question informs models of resource allocation in working memory, as we discuss below. We found that an item’s predicted utility modulates its fidelity in WM, even when it is the only item being maintained and so there is no concurrent load or difference in relative utility from other items in memory. As discussed below, this result presents a subtle but fundamental distinction from prior conceptualizations of resource allocation in WM.

We parametrically manipulated predicted utility such that an item encountered at encoding had a likelihood of 0%, 50%, or 100% of being tested on that trial. Using a generative modeling method of analyzing multivoxel fMRI activity from visual cortex, we estimated orientation probability distributions during the maintenance delay period following the presentation of the first grating. As such, the load was matched at one item for the medium- and high-PU conditions. Thus, any differences in the measures of representational fidelity between these conditions could not be attributed to differences in load or the relative utility among items in memory, but rather what the participant knew about the predicted utility of that first item.

We hypothesized that predicted utility would influence the fidelity of orientation representations in visual cortex during WM. We operationalized fidelity using entropy and posterior SD as these measure the uncertainty about the remembered orientation. Consistent with our hypothesis, the entropy computed over orientation probabilities decreased parametrically with increasing predicted utility. Thus, even when there are no other items in memory to share resources with a higher-priority item, items are nonetheless maintained with a fidelity accorded by their predicted utility.

It is important to emphasize that these differences in predicted utility condition were not dependent on comparison of the high- and medium-PU conditions to no-PU condition only. The no-PU condition is arguably a case where participants may ignore the item entirely, and not encode it in working memory, and on its own, its difference from the other two conditions might be one of state (i.e., remember or not) rather than degree. Further, there is already evidence of differences in memory fidelity and performance between known targets and distractors, which is equivalent to our high-PU and no-PU conditions in terms of predicted utility (McNab & Klingberg, 2008). We replicate those differences here, but extend them by estimating the fidelity of the remembered representation and probing a graded response with inclusion of the medium-PU condition. The crucial difference between high- and medium-PU items was evident in entropy and posterior SD as we hypothesized, and notably, also in the probability of the presented orientation. Furthermore, we reproduced this difference with our own dataset using the psychologically equivalent trials with future distractors first instead of the future targets in the medium-PU condition.

The observation that the fidelity of representations in working memory are modulated (a) separately from load and (b) in a graded fashion based on predicted utility has important implications for theories of resource allocation in working memory. First, this observation is inconsistent with passive mechanisms of resource allocation, such as normalization of neural activity patterns over maintained items in memory (Bays, 2014) or the limits of spacing of items in a phase space of a particular oscillatory frequency (Lisman & Idiart, 1995; Luck & Vogel, 2013; Raffone & Wolters, 2001). In this experiment, there are no other items competing for resources with the first item in the high- and medium-PU conditions. Rather, these data suggest that there must be an active resource allocation mechanism – like a WM gate or an attentional filter on WM input – capable of pre-allocating resources based on the item’s own predicted utility and without regard to its relative utility to other items currently in memory.

One possibility that we cannot rule out in the present design is that resources are allocated based on expectations about future load rather than expected utility. Though participants did not know whether the first item in the context-last condition would be a target or distractor, they were aware that a second item to remember would appear after this item. So, one possibility is that they somehow allocated resources in expectation of two items rather than allocating fewer resources to this one item based on its medium level of predicted utility. Though this is a plausible account, we do not know of models that propose such a mechanism presently. And, regardless, this account of working memory still requires an active mechanism that can proactively assign resources to items, though here based on expected load rather than expected utility. This will be a point to resolve in future research.

This possibility notwithstanding, this experiment provides strong evidence that the quality of working memory representations is determined not only by current load demands but also by the utility of the items themselves for later behavior. In other words, the brain allocates resources in accord with its predictions about how useful maintenance of that item should be.

These results align well with the recently proposed resource-rational theory of WM (Ronald van den Berg & Ma, 2018). This theory says that WM resources are not fixed. Instead of allocating resources based on set size (i.e., allocating less resource towards each item when there are more items to remember), we instead optimize resource allocation by weighing the tradeoff between expected behavioral cost and the cost of expending neural resources on encoding. That is, we want to perform well but we want to use as few neural resources as possible to do so. Their model shows that the set size effect (i.e., the finding that precision decreases with set size) is mediated by probe probability and not number of items to be remembered. That is, more resource is devoted to items when the likelihood of those items being probed is higher, but when more items are presented with equal probe probabilities, the relevance of any of those items being probed goes down (Emrich et al., 2017), and thus it is less worth it to expend resources on encoding those items. This is relevant to the current study because the context-first and context-last conditions have equal set sizes but unequal probe probabilities (1 in the context-first condition and ½ in the context-last condition). Thus, one explanation for why we see precision differences across conditions may be because there is a higher cost to storing items with high precision.

As already noted, these results also favor models in which memory capacity is managed by an active mechanism that takes account of predicted utility of maintenance. This fits well with gated WM models that include mechanisms for connecting value to the control of memory. Several models of gated memory have been proposed (Badre & Frank, 2012; Barak & Tsodyks, 2014; Braver & Cohen, 2000; Conde-Sousa & Aguiar, 2013; Dipoppa & Gutkin, 2013; Edin et al., 2009; Frank et al., 2001; Frank & Badre, 2012; Hazy et al., 2007; Mante et al., 2013; Murray et al., 2017; O’Reilly & Frank, 2006; Santos et al., 2012; Strock et al., 2020; X.-J. Wang et al., 2004; Zhu et al., 2020). These gated memory models commonly detail mechanisms by which items encountered at one time point can be placed in a state such that they are maintained robustly against distractors and so are accessible in a high-fidelity form at a later time point when needed.

Importantly, a subset of these models emphasize the action of the neurotransmitter dopamine to explain how WM might be modulated based on value (Badre & Frank, 2012; Braver & Cohen, 2000; Frank et al., 2001; Frank & Badre, 2012; Frank & O’Reilly, 2006; Hazy et al., 2007). Evidence in both animals and humans has linked cortical dopamine to improved accuracy in WM tasks and maintenance that is more robust to distraction (Durstewitz & Seamans, 2002; Frank & Fossella, 2011; Furman et al., 2020; Wiecki & Frank, 2013). In essence, dopamine increases the signal-to-noise ratio (SNR) in cortex, which may be in line with our observation of progressively reduced entropy for higher predicted utility items. Further, gating mechanisms, such as through cortico-striatal loops, that incorporate DA as a modulator can leverage DA as a learning signal to influence what items are gated based on their predicted future value.

An alternative mechanism to gating that could also potentially account for the observed effects of predicted utility might involve attentional modulation. In particular, predicted utility might lead to an attentional bias for lines oriented in a certain range on high-PU context-first trials, which in turn could enhance or suppress responsibility to orientations inside or outside that range respectively, relative to medium-PU context-last trials. That is, the initially presented context in high-PU trials could allow for feature-based encoding earlier in the information processing pipeline, relative to medium-PU trials which do not utilize a pre-cue. In this case, the mechanism for modulating WM fidelity as a function of predicted utility would be an attentional filter on WM input.

We attempted to distinguish between an attentional account and a gating account of our data by analyzing TR windows later in the first-grating delay window. We reasoned that an attentional cueing effect would be evident early in the WM delay, even at the point of stimulus processing and would carry through the delay. By contrast, the benefits of gating could emerge later, particularly when noise has greater opportunity to disrupt maintenance of items that have not been gated.

We found that the probability of the presented/remembered orientation was highest for the latest time window (10-14 seconds after delay onset) compared to the earlier time windows (6-10 seconds and 8-12 seconds after delay onset). These observations suggest that the representation strength of items being held in WM are being modulated by predicted utility later in the information processing pipeline, even at the point where they must be held robust to interference from the second item. In our view, this is more in line with a WM gating than attentional pre-cueing account.

We note that this late modulation may be consistent with observations from retrocueing studies finding that post-encoding prioritization formats items in memory for their use, linking them to upcoming actions (Myers et al., 2017; Nobre & Stokes, 2019; Olivers & Roelfsema, 2020; Orhan & Ma, 2019; Souza & Oberauer, 2016b). It is also consistent with recent observations we have made using EEG wherein action plans held in WM are prioritized late, even when their priority is pre-cued (Kikumoto and Badre, 2022, biorxiv).

Though we find our results fit better with a gating account than an attentional one, there is also a modulation by PU earlier in the delay. Since our task was not designed with this time window comparison in mind, the observation of late modulation should be replicated before definitely ruling out an attentional contribution or to assess the contribution of each mechanism to PU modulation of working memory. Indeed, a task design that manipulates attention at the cue stage relative to within the maintenance delay may be better able to establish whether gating or attention (or another mechanism) best explain how predicted utility modulates fidelity.

In summary, we provide evidence that predicted utility modulates the fidelity of working memory representations in the brain. This effect is independent of load or the relative utility of an item to other items in memory, and as such, it favors working memory models that employ value-based selective gating mechanisms to input items in a graded way for maintenance in working memory.

## Acknowledgements

We thank Lakshmi Govindarajan and Apoorva Bhandari for their help and input with experiment design and analysis, Emily Chicklis, Adriane Spiro, and Johanny Castillo for their assistance with data collection, and members of the Badre lab for their help and input in this study. This work was supported a MURI award from the Office of Naval Research (N00014-16-1-2832), an R01 from NIMH (MH111737), and an award from the James S. McDonnell Foundation. We are grateful to members of the Badre Lab for their help and input on this study.

## Notes

### Competing Interest Statement

The authors have declared no competing interest.

### Summary of Updates

Among our revisions, we have incorporated discussion of an attentional model as a plausible explanation for our findings that can be distinguished from a gating model, and now include an additional time window analysis in our results section in an attempt to further differentiate these two accounts. We also now include a trial-level analysis that differentiates between whether posteriors are wider at the single-trial level, or narrower but more inaccurate. In addition to entropy, we also now include posterior standard deviation as an additional measure of uncertainty at both the trial-averaged and single-trial levels, as posterior SD is computed based on the variance in angles about the mean direction.

